# Conserved collateral susceptibility networks in diverse clinical strains of *Escherichia coli*

**DOI:** 10.1101/248872

**Authors:** Nicole L. Podnecky, Elizabeth G. A. Fredheim, Julia Kloos, Vidar Sørum, Raul Primicerio, Adam P. Roberts, Daniel E. Rozen, Ørjan Samuelsen, Pål J. Johnsen

**Affiliations:** Department of Pharmacy, Faculty of Health Sciences, UiT The Arctic University of Norway, Tromsø, Norway; Department of Parasitology, Liverpool School of Tropical Medicine, Liverpool, UK; Research Centre for Drugs and Diagnostics, Liverpool School of Tropical Medicine, Liverpool, UK; Institute of Biology, Leiden University, Leiden, The Netherlands; Norwegian National Advisory Unit on Detection of Antimicrobial Resistance, Department of Microbiology and Infection Control, University Hospital of North Norway, Tromsø, Norway.

**Keywords:** Collateral sensitivity, cross-resistance, antimicrobial resistance, fitness cost of resistance, selection inversion

## Abstract

There is urgent need to develop novel treatment strategies to reduce antimicrobial resistance. Collateral sensitivity (CS), where resistance to one antimicrobial increases susceptibility to other drugs, is a uniquely promising strategy that enables selection against resistance during treatment. However, using CS-informed therapy depends on conserved CS networks across genetically diverse bacterial strains. We examined CS conservation in 10 clinical strains of *E. coli* resistant to four clinically relevant antibiotics. Collateral susceptibilities of these 40 resistant mutants were then determined against a panel of 16 antibiotics. Multivariate statistical analyses demonstrate that resistance mechanisms, in particular efflux-related mutations, as well as relative fitness were principal contributors to collateral changes. Moreover, collateral responses shifted the mutant selection window suggesting that CS-informed therapies could affect evolutionary trajectories of antimicrobial resistance. Our data allow optimism for CS-informed therapy and further suggest that early detection of resistance mechanisms is important to accurately predict collateral antimicrobial responses.

---

The evolution and increasing frequency of antimicrobial resistance (AMR) in bacterial populations is driven by the consumption and misuse of antimicrobials in human medicine, agriculture, and the environment (1-3). Historically, the threat of AMR was overcome by using novel antimicrobials with unique drug targets. However, the discovery rate of new antimicrobial agents has dwindled (4-6) and severely lags behind the rate of AMR evolution. While concerted scientific, corporate and political focus is needed to recover antimicrobial pipelines (7-9), there is an urgent need for alternative strategies that prolong the efficacy of existing antimicrobials and prevent or slow the emergence, spread and persistence of AMR. Current global efforts to improve antimicrobial stewardship largely focus on awareness and reducing overall consumption (8, 10-12). While these efforts will affect the evolution and spread of AMR, mounting evidence suggests that these changes alone will not lead to large-scale reductions in the occurrence of AMR (13-17).

Several recent studies have examined novel treatment strategies using multiple antimicrobials that could reduce the rate of resistance emergence and even reverse pre-existing AMR. These approaches, collectively termed “selection inversion” strategies, refer to cases where resistance becomes costly in the presence of other antimicrobial agents (18). Among the most promising of these strategies are those based on a phenomenon first reported in 1952 termed “collateral sensitivity” (CS), where resistance to one antimicrobial simultaneously increases the susceptibility to another (19). CS and its inverse, cross-resistance (CR), have been demonstrated for several bacterial species and across different classes of antimicrobials (20-26). These results have formed the basis of proposed CS-informed antimicrobial strategies that mix drug pairs (21, 27) or alter temporal administration, e.g. drug cycling (20, 28). The notion behind these strategies is that they would force bacteria to evolve AMR along a predictable trajectory that results in CS; this predictability can then be exploited to ultimately reverse resistance and prevent the fixation of AMR and multi-drug resistance development at the population level of bacterial communities.

Initial *in vitro* experiments support using CS-based strategies to re-sensitize resistant strains (20) and reduce rates of resistance development (28); however, the broader application of this principle depends on predictable bacterial responses during antimicrobial therapy. This predictability must be general for a given drug class and should not vary across strains of the same species. To date, most studies of CS and CR have focused on describing these responses as collateral networks using resistant mutants derived from single laboratory adapted strains and few exemplary clinical isolates. Two studies on *Pseudomonas aeruginosa* have further investigated CS in collections of clinical isolates (29, 30). However, these studies lack either proper baseline controls (29) or sufficient genetic diversity (30). As valuable as previous studies have been, the responses of single strains (laboratory or clinical) may not be representative of CS and CR evolution in other strains. To address this limitation, we focused on understanding collateral networks in clinical urinary tract infection isolates of *Escherichia coli* with selected resistance to drugs widely used for the treatment of urinary tract infections; ciprofloxacin (CIP), trimethoprim (TMP), nitrofurantoin (NIT), or mecillinam (MEC).

We investigated the collateral networks to 16 antimicrobials from diverse drug classes in 10 genetically diverse clinical strains (corresponding to 40 laboratory-generated AMR mutants) to assess the factors contributing to collateral responses (both CS and CR). This approach allowed us to identify variation in the sign and magnitude of collateral responses and identify mechanisms of CS and CR that are preserved in different genetic backgrounds. By using multivariate statistical modeling of experimental data, we show that resistance mutations, particularly those affecting efflux pumps, and the relative fitness of resistant isolates are more important determinants of the structure of collateral networks than genetic background. Our results support the idea that collateral responses may be predictable.

## Results

### Collateral responses are frequent and vary both between and across AMR groups

Starting from 10 pan-susceptible urinary tract clinical *E. coli* isolates (**Fig. S1**) (31), a single AMR mutant was generated to each of four individual antimicrobials used for urinary tract infections treatment, CIP, TMP, NIT, and MEC. Here we define ‘AMR group’ as the collection of mutants from the 10 different genetic backgrounds that were selected for resistance to the same antimicrobial. In total 40 AMR mutants were generated that had resistance levels above clinical breakpoints, as determined by antimicrobial susceptibility testing using both gradient strip diffusion (**Table S1**) and inhibitory concentration 90% (IC_90_; (20)) testing (Table 1). Changes in the IC_90_ of AMR mutants from each respective wild-type (WT) strain (**Fig. S2**) were compared for 16 antimicrobials (Table 2). The two methods are correlated but IC_90_ measurements allow for more robust detections of small relative differences in susceptibility (32, 33). Overall, collateral responses were observed in 40.5% (243/600) of possible instances (**Table S2**); of these, roughly half, 47% (115/243), were associated with only a 1.5 and 2-fold change in IC_90_. Such small changes would not be observed by typical 2-fold antimicrobial susceptibility testing methods frequently used in clinical laboratories.

**Table 1:**
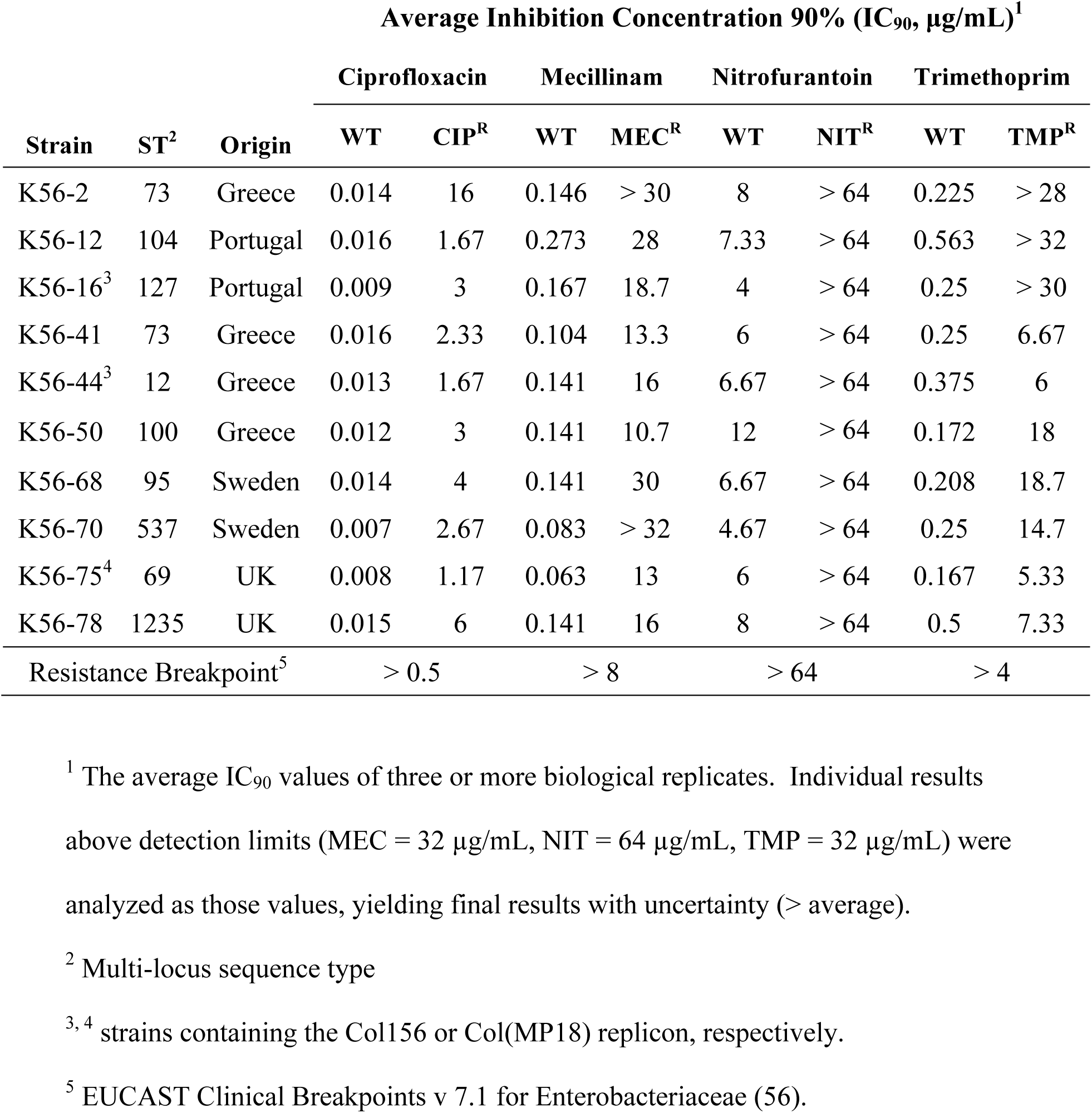
Description of *Escherichia coli* strains used in the study and susceptibility changes following antimicrobial selection *in vitro*.

**Table 2:** List of antimicrobials used in this study.

| Antimicrobial<sup>1</sup> | Abbreviation | Drug Class | Drug Target(s) | Solvent |
| --- | --- | --- | --- | --- |
| Amoxicillin | AMX | β-lactam (Penicillin) | Cell wall synthesis | Phosphate buffer<sup>3</sup> |
| Azithromycin | AZT | Macrolide | Protein synthesis (50S) | $\geq 95\%$ Ethanol |
| Ceftazidime | CAZ | β-lactam (Cephalosporin) | Cell wall synthesis | Water + 10% (w/w) Na<sub>2</sub>CO<sub>3</sub> |
| Chloramphenicol | CHL | Amphenicol | Protein synthesis (50S) | $\geq 95\%$ Ethanol |
| Ciprofloxacin | CIP | Fluoroquinolone | DNA replication, cell division | 0.1N HCl |
| Colistin | COL | Polymyxin | Cell wall & cell membrane | Water |
| Ertapenem | ETP | β-lactam (Carbapenem) | Cell wall synthesis | Water |
| Fosfomycin | FOS | Phosphonic | Cell wall synthesis (MurA) | Water |
| Gentamicin | GEN | Aminoglycoside | Protein synthesis (30S) | Water |
| Mecillinam | MEC | β-lactam (Penicillin) | Cell wall synthesis (PBP2) | Water |
| Nitrofurantoin | NIT | Nitrofuran | Multiple<sup>4</sup> | Dimethyl sulfoxide |
| Trimethoprim | TMP | Antifolate | Folate synthesis (FolA) | Dimethyl sulfoxide |
| Sulfamethoxazole | SMX | Antifolate | Folate synthesis (FolP) | Dimethyl sulfoxide |
| TMP+SMX (1:19) | SXT | Antifolate | Folate synthesis (FolA+FolP) | Dimethyl sulfoxide |
| Temocillin | TEM | β-lactam (Penicillin) | Cell wall synthesis | Water |
| Tetracycline | TET | Tetracycline | Protein synthesis (30S) | Water |
| Tigecycline | TGC | Tetracycline | Protein synthesis (30S) | Water |

<sup>1</sup> When available final antimicrobial concentration was determined using manufacturer-provided or calculated drug potencies, otherwise potency was assumed to be 100%. Aliquots were stored at -20°C or -80°C in single use vials. All antimicrobials and chemical solvents were obtained from Sigma-Aldrich (St. Louis, MO, USA) with the exception of CIP (Biochemika, now Sigma-Aldrich) and TEM (Negaban®).
<sup>3</sup> 0.1 mol/L, pH 6.0 phosphate buffer supplemented with 6.5% (v/v) 1M NaOH (sodium hydroxide).
<sup>4</sup> Nitrofurantoin is thought to target macromolecules including DNA and ribosomal proteins, affecting multiple cellular processes including protein, DNA, RNA and cell wall synthesis.

Overall CR was more frequent than CS, 151 versus 92 instances (**Table S2**), and collateral networks varied considerably between AMR groups. We observed 20 cases of conserved collateral responses (Fig. 1), where CR or CS to a specific antimicrobial was found in ≥ 50% of the mutants within an AMR group, defined as CR50 or CS50 respectively. This indicates that some collateral responses are likely general, irrespective of genetic background. For each CS_50_ and CR_50_ observation, IC_90_ results were further assessed by generating dose response curves of representative strain:drug combinations (**Fig. S3**). Inhibition of growth was shown to vary across antimicrobial concentrations between AMR mutants and respective WT strains, confirming the changes in antimicrobial susceptibility determined by the IC_90_ assays. While conserved collateral responses were observed, there was variability across the remaining genetic backgrounds.

**Figure 1.**
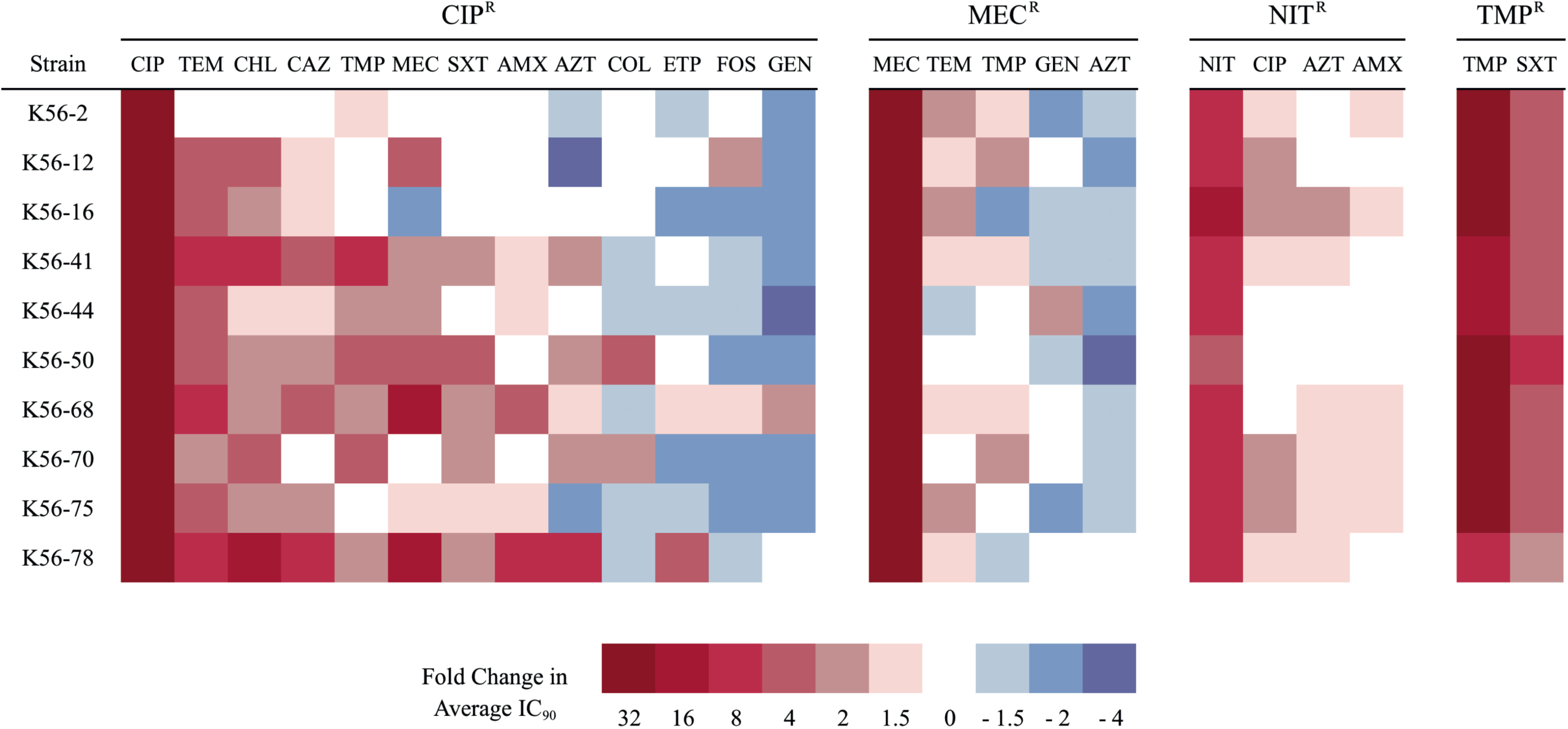
Conserved collateral effects observed in AMR mutants of different AMR groups. Relative change in antimicrobial susceptibility was determined by comparing average IC_90_ values of AMR mutants to their respective WT strain. Collateral changes that were found in ≥ 50% of the strains are displayed. Antimicrobials are ordered by most frequent CR (left) to most frequent CS (right) for each AMR group. The only instances where CR was present in 100% of the strains of an AMR group were linked to the drug used for selection, including TMP^R^ strains with CR to SXT (a combination of TMP and SMX).

During the selection of AMR mutants, we often observed conspicuous changes to colony size for all AMR groups, suggesting that resistant mutants had reduced bacterial fitness. To test this, we measured the growth rates of AMR mutants and compared these to the respective WT strains (**Fig. S4**). In general, CIP^R^ and MEC^R^ mutants displayed severely reduced growth rates suggesting high costs of resistance. The cost of CIP^R^ and MEC^R^ mutants, measured as relative growth rate, was significantly different from 1 and varied between 0.34 and 0.75 with a mean of 0.53 for all CIP^R^ mutants, whereas the cost of MEC^R^ mutants varied between 0.49 and 0.79 with a mean of 0.64. NIT^R^ and TMP^R^ mutants displayed lower levels of reduced fitness and several resistant mutants harbored apparent cost-free resistance mutations (**Fig. S4**). Only two of ten NIT^R^ mutants and four of ten TMP^R^ mutants displayed a significant cost of resistance. Relative growth rates varied between 0.93 and 1.05 and 0.68 and 1.07 with averages of 0.99 and 0.94 for NIT^R^ and TMP^R^ mutants, respectively.

### Conserved collateral responses were primarily found in CIP^R^ mutants

Nearly half (108/243, 44%) of the observed collateral responses were in CIP^R^ mutants, while the remaining 135 were distributed between the other three AMR groups (**Table S2**). Within the CIP^R^ group, the majority of collateral responses were CR (70/108, 64.8%). Additionally, CS responses found for the CIP^R^ mutants were the most conserved in our dataset, with CS to GEN occurring in 8 of 10 strains and CS to FOS in 7 of 10 strains. GEN and other aminoglycosides are important for the treatment of a wide range of hospital infections (34), while FOS is primarily used for treatment of uncomplicated urinary tract infections (35, 36). The CIP^R^ mutants were also unique in the magnitude of observed changes, with cases of CR close to 30-fold and CS as high as 6-fold changes in IC90 (**Fig. S2**). Only CIP^R^ isolates displayed potential clinically relevant CR, where six strains had IC_90_ values above the clinical breakpoint for resistance to CHL and a single strain (K56-68 CIP^R^) was above the AMX breakpoint.

### Characterization of AMR mutants

We hypothesized that CS and CR variation in and between the AMR groups could be attributed to mutations causing resistance in each strain. Using whole genome sequencing we identified a total of 150 mutations in the AMR mutants (**Table S3**). Of these, 88 mutations affect previously described or putative AMR associated genes, gene regions, or pathways (**Table S3**). The remaining mutations were found in other cellular processes with no known association to antimicrobial resistance (*e.g.* metabolic pathways and virulence factors), such as mutation to the FimE regulator of FimA that was frequently observed in MEC^R^ mutants (**Table S3**). Aside from FimE, we did not observe similar mutations in non-AMR associated regions across strains of the same AMR group (parallel evolution), suggesting that such mutations had limited, if any effect on collateral responses in this study. For each of the 40 AMR mutants at least one putative resistance mechanism was identified, including mutations to previously described antimicrobial drug targets and promoters of drug targets, drug modifying (activating) enzymes, regulators of efflux pumps, RNA polymerases and mutations to other metabolic and biochemical processes that may contribute to resistance (Table 3). Briefly, all but one CIP^R^ mutant contained mutations in *gyrA* and efflux regulatory genes and gene-regions likely affecting efflux expression (*acrAB* and/or *mdtK*), while one strain had only drug target mutations and displayed the well-described GyrA (S83L) and ParC (G78D) mutation combination. Both efflux and drug target mutations are frequently found in surveys of clinical isolates (37-40). NIT^R^ mutants had mutations in one or both nitro-reductases (*nfsA*, *nfsB*) and the majority of strains had additional mutations in *mprA* that encodes an efflux regulator of EmrAB-TolC pump expression. TMP^R^ mutants contained mutations either in *folA* and/or its promoter or genetic amplification of a large genetic region containing *folA.* MEC^R^ mutants are unique in that they were evolved as single step mutants, where a single mutation could confer clinically relevant resistance to MEC. Resistance development for the remaining three drugs required several steps, as multiple mutations were required for resistance above clinical breakpoints. In total 12 different mutations in genes and/or cellular processes previously linked to MEC^R^ were identified in the MEC^R^ group (41).

**Table 3:** The number of AMR mutants with resistance-associated mutations.

| AMR Mechanism |  | CIP<sup>R</sup> | MEC<sup>R</sup> | NIT<sup>R</sup> | TMP<sup>R</sup> |
| --- | --- | --- | --- | --- | --- |
| Drug Target | Modification | 10<sup>1</sup> |  |  | 6 |
|  | Overproduction |  |  |  | 6 |
| Drug Activation | Nitroreductase disruption |  |  | 10 |  |
| Drug Uptake | Porin mutation | 1 |  |  |  |
| Efflux | AcrAB-TolC | 7 |  | 1 |  |
|  | MdtK | 9 |  | 1 |  |
|  | MdfA |  |  | 1 |  |
|  | EmrAB-TolC |  |  | 7 |  |
|  | ABC transport |  | 1 |  |  |
| ppGpp synthesis<br>(stringent response<br>activation) | Stringent Response |  | 4 |  |  |
|  | tRNA synthesis |  | 4 |  |  |
|  | tRNA processing |  | 1 |  |  |
|  | Cellular metabolism |  | 3 |  |  |

<sup>1</sup> All CIP<sup>R</sup> mutants contained one mutation in the *gyrA* gene, except K56-2 CIP<sup>R</sup> mutant that contained two mutations in *gyrA* and a mutation in *parC*.

Our data suggest that the few collateral responses in TMP^R^ mutants are attributable to a specific mechanism of resistance affecting a single unique drug target (*i.e.* overexpression/alteration of FolA). Similarly, the CIP^R^ group displayed a clear trend where conserved CR responses were strongly linked to mutations in efflux regulatory regions. In the remaining AMR groups we observed heterogeneous collateral responses.

The WT genomes were evaluated for the presence of genetic elements linked to AMR. We detected only one acquired genetic element, *sul2* (linked to sulfonamide resistance) in K56-44, and two point mutations, PmrB V161G in K56-50 and K56-70 and ParE D475E in K56-78, that are linked to COL and quinolone (CIP) resistance, respectively (**Table S3E**). It is unclear what, if any, effect these resistance determinants have in these WT strains since all were phenotypically pan-susceptible (**Fig. S1**). However, the ParE mutation could contribute to CIP^R^ levels in K56-78, as the K56-78 WT was among the higher WT values for CIP (0.015 μg/mL) (**Fig. S1**) and K56-78 CIP^R^ was among the highest IC_90_ values of the CIP^R^ mutants (Table 1).

### Multivariate statistical analyses suggest that efflux-related mutations and relative fitness are significant contributors to collateral responses

Multivariate statistical approaches were used to investigate the extent to which genetic (strain) background, AMR group, the putative mechanism of resistance in particular efflux-related mutations (**Table S3**), growth rate and the fitness cost of resistance explain the total variation in collateral responses. All the above factors were investigated individually and the related models are found in the supplementary material (**Fig. S5A**). Throughout the remaining analyses we focus mainly on efflux-related mutations rather than AMR group to explicitly address putative mechanisms of resistance and on relative fitness rather than the growth rate. We estimated several models with individual, or a combination of, factors to assess their effect size and significance given some level of co-linearity between fitness and efflux type (Fig 2, **Fig. S5C-F**). A model including strain background, relative fitness and efflux-related mutations as factors explained 62.53% of the total variation in IC_90_ values (Fig. 2A, **Table S4**). In this three-factor model there was clear separation of the mutants by AMR group. The CIP^R^ mutants showed strong CR towards TEM, CHL, CAZ and AMX separating this AMR group from the others along the first ordination axis. Along the second ordination axis, MEC^R^ isolates were distinct, had CR to TEM, and were more likely to have CS towards drugs such as AZT and CHL (Fig. 2A). Both efflux type and relative fitness were significant predictors when tested alone and in combination (**Table S4**). The model (Fig. 2A) also revealed that strain background had a limited, non-significant contribution. Even when modeled alone (**Fig. S5A**), strain background only accounted for 6.53% of the variation and was non-significant (**Table S4**).

**Figure 2.**
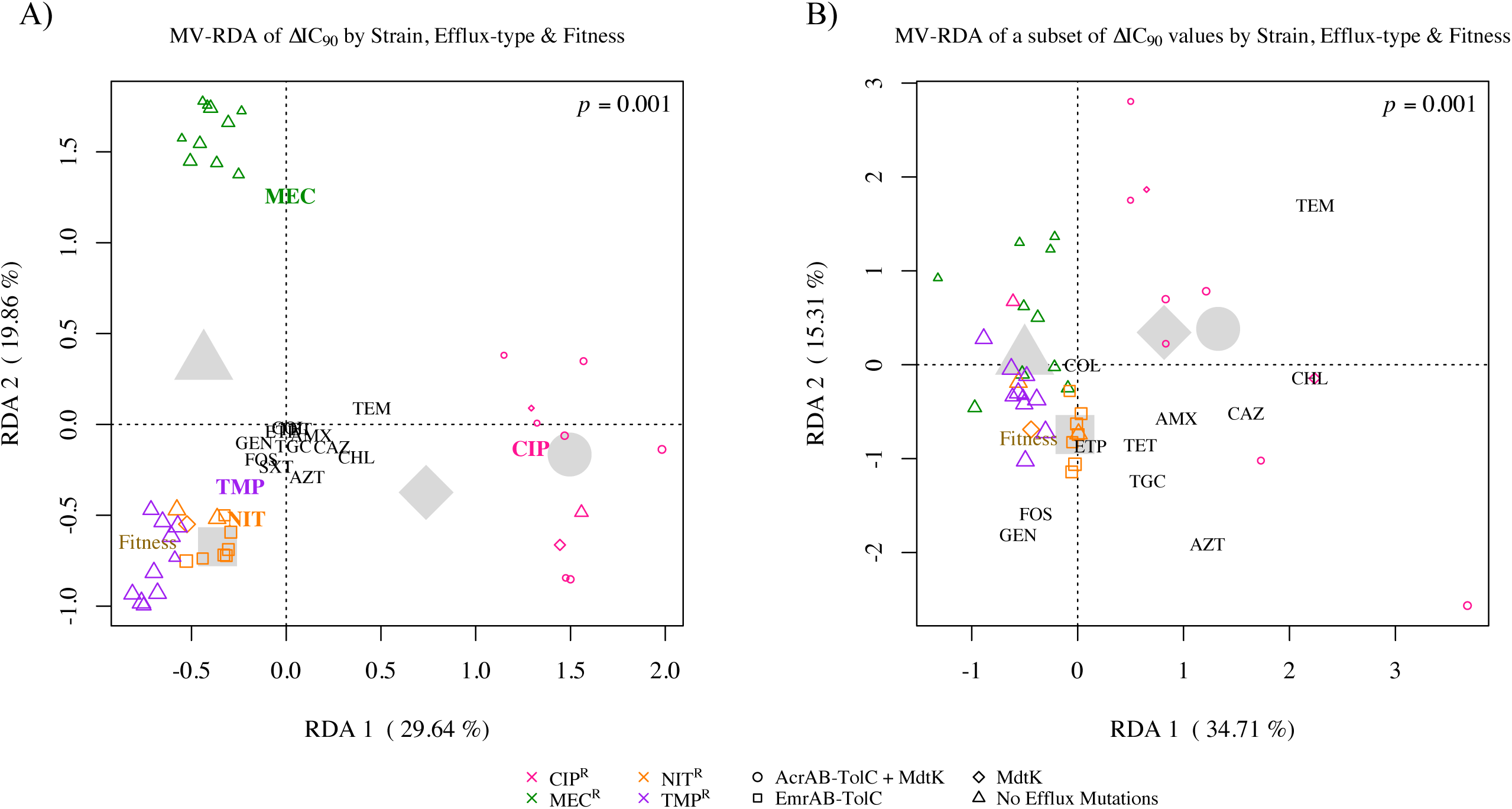
Results of multivariate statistical modeling. Graphical representations of redundancy analysis (RDA, triplot) results relating strain background, presence of efflux-related mutations and relative fitness to the observed changes in IC_90_ between AMR mutants and the respective WT strain for 16 antimicrobials tested (**A**) and a subset of the antimicrobials excluding CIP, MEC, NIT, TMP and SXT (**B**). The first and second RDA axes shown display most of the explained variation in IC_90_ changes. The weighted average of each AMR mutant is plotted as a single colored symbol, where color indicates the AMR group (CIP^R^ – pink, MEC^R^ – green, NIT^R^ – gold, TMP^R^ – purple), shape the assigned efflux type (circle – AcrAB-TolC + MdtK, square – EmrAB-TolC, diamond – MdtK, triangle – no efflux-related mutations), and symbol size is proportional to relative fitness, where smaller size indicates a greater reduction in growth rate compared to the WT. Antimicrobial drug names indicate the tip of vectors that pass through the origin in the direction of increasing IC_90_ fold change or CR (direction of steepest ascent). These vectors can be used to interpret the change in IC_90_ for the antimicrobials shown, e.g. there was little change in the IC_90_ of drugs centered near the origin, such as COL in **B**. The vector tip of relative fitness (brown) is also shown. Large grey symbols show the centroids (average effect) for all AMR mutants within a given efflux group (shape). The majority of explained variation is driven by primary resistances (**A**), where CIP^R^ mutants (pink) cluster away from the other three AMR groups along the CIP vector, indicating higher resistance to CIP. CIP^R^ mutants are likely to show CR to CHL, CAZ, TEM and AZT, but sensitivity to GEN, FOS and TMP. MEC^R^ isolates primarily display low-level CS to most antimicrobials tested. TMP^R^ and NIT^R^ groups cluster together with relatively few collateral effects. The analysis of the subset (**B**) shows patterns consistent with the full model, but with less clustering by AMR group. However, in the redundancy analysis on a subset of the IC_90_ fold changes (**B**) there is far less clustering of the AMR mutants by AMR group (color).

We initially hypothesized that genetic background would significantly affect collateral responses. Our data suggest that it does not. Arguably, the inclusion of IC_90_ data from the drugs to which primary resistance was selected for could confound the analysis, despite our efforts to minimize these effects using log transformed data. We used the same approaches to assess a subset of collateral responses, excluding data for all of the 40 AMR mutants to five antimicrobials containing the drugs used for selection (CIP, MEC, NIT, TMP) and SXT. Within the subset model, patterns consistent with the full model were observed, but with a lower degree of clustering by AMR group (Fig. 2B). For example, the K56-2 CIP^R^ mutant is now co-localized with the MEC^R^ isolates, indicating that this isolate is distinct from other CIP^R^ mutants, which still showed strong tendencies of CR to TEM, CHL, CAZ and AMX. Despite these changes, efflux type and fitness were still significant predictors of collateral networks, whereas strain background remained non-significant in models, both in combination and alone (**Fig. S5B**, **Table S4**). Mutations in efflux-related genes and gene regulators were the strongest predictor of collateral responses tested, explaining over 33% of the variation in the subset. Fitness alone also had significant predictive value, but to a lesser extent (17% variation explained). It is important to note that we observed correlation between efflux mutations and relative fitness that is likely explained by reduced fitness resulting from the cost of overexpression of efflux pump(s) (38).

To investigate the influence of resistance mechanism on IC_90_ variation at a higher resolution, we modeled each AMR group separately relating the putative resistance mechanism and fitness separately and in combination (**Fig. S5**). However potentially due to a lower number of samples within AMR groups and more detailed classification of the resistance mechanism, these factors had varying degrees of contribution. For TMP^R^ and CIP^R^ AMR groups, resistance mechanism was non-significant, but it was a significant factor for NIT^R^ and MEC^R^ mutants. Fitness was a significant factor of variation only for the MEC^R^ group and similarly, combination models using both resistance mechanism and fitness were non-significant for all AMR groups, with the exception of the MEC^R^ group.

### Collateral responses shift the mutation selection window

The mutant selection window (MSW) can be defined as the concentration space between the lowest antimicrobial concentration that selects for and enriches resistant mutants (42) and the concentration that prevents the emergence of first step resistant mutants, the mutation prevention concentration (MPC) (43, 44). In theory, if drug concentrations remain above the MPC during treatment AMR is less likely to evolve (43, 44). It was recently demonstrated in *E. coli* MG1655 that changes in MPC correlated with collateral responses in AMR mutants (20). We determined the MPC for 17 strain:drug combinations that exemplified the conserved collateral responses (Fig. 1). The MPC for each AMR mutant and its respective WT were compared (Fig. 3). In 12/17 (70.6%) the change in MPC was consistent with the sign of collateral responses in IC_90_. This demonstrates that even small CS/CR changes can affect the MSW, correspondingly shifting it down or up. However, in 4/17 (23.5%) the MPC displayed no change between the WT and AMR mutant. This was observed when testing the MPC for MEC, TMP and AZT, though we speculate that increasing the precision of the MPC assay (as was done with IC_90_ testing) might negate these discrepancies. Changes in MPC results with AZT were inconsistent with the change in IC_90_ for a CIP^R^ mutant and the MEC^R^ and NIT^R^ mutants, which displayed a decreased MPC instead of an expected increase or no change, respectively.

**Figure 3.**
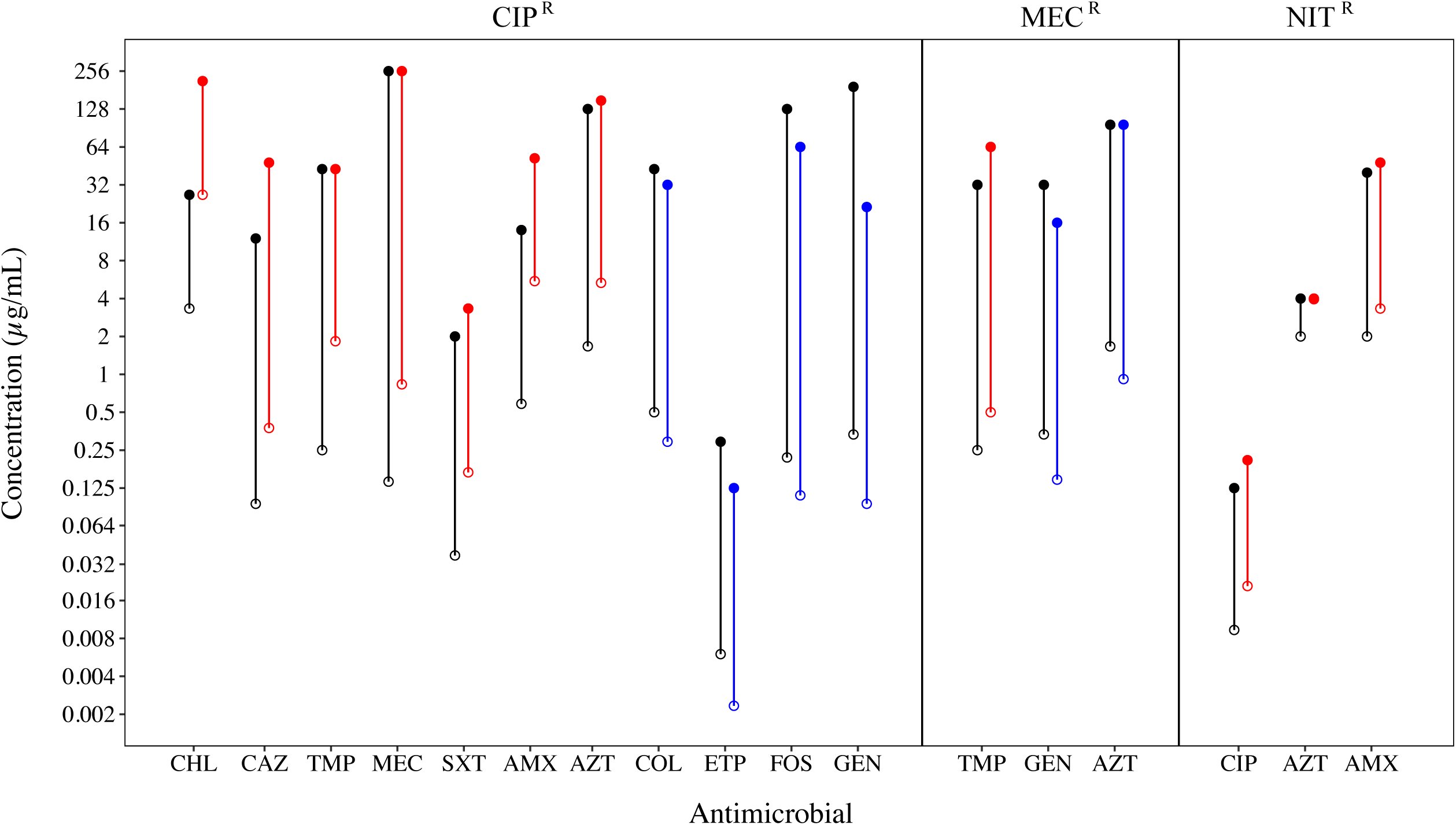
Summary of changes in IC_90_ and MPC results for representative strain:drug combinations. The average IC_90_ (open circles) and average mutation prevention concentration (MPC; filled circles) were determined and compared between AMR mutants (colored) with collateral responses, either CS (blue) or CR (red), and their respective wild-type strain (WT; black) in strain:drug combinations representing conserved collateral responses. The range between the IC_90_ and MPC was considered the mutation selection window (MSW; lines). K56-16 NIT^R^ had equivalent IC_90_ and MPC values for AZT, thus no MSW was reported. Generally, changes in MPC values reflected observed IC_90_ changes, shifting the MSW upwards or downwards accordingly. In 8/10 tested combinations an increase in IC_90_ value (CR) from WT to AMR mutant correlated with at least a small increased MPC, with the remaining combinations showing no change in MPC value between the WT and AMR mutant. Similarly, decreased IC_90_ values (CS) correlated with decreased MPCs (5/7).

## Discussion

Here we present, to our knowledge, the first in depth study to identify conserved collateral responses in antimicrobial susceptibility across genetically diverse clinical *E. coli* strains following antimicrobial resistance development. Our findings are relevant beyond urinary tract infections because uropathogenic *E. coli* are shown to also stably colonize the bladder and gut (45), and to cause bloodstream infections (46). Our data show that CS and CR are pervasive in clinical *E. coli* strains, consistent with earlier results based on laboratory-adapted strains of various species (20-22, 24, 29, 47) and a limited number of clinical isolates (20, 29). Resistance to CIP resulted in a greater number of collateral responses than resistance to MEC, NIT, or TMP. CR was much more prevalent than CS, and the magnitude of the collateral responses were most often small, consistent with other reports (20-22). Overall we observed that collateral responses varied substantially by AMR group, but variation was also observed within AMR groups.

Using CS_50_ and CR_50_ thresholds to identify conserved responses, we found that conserved CR was more than twice as common as conserved CS. Whereas many of the conserved collateral responses identified in this study support the findings in previous work using single laboratory-adapted strains, we observed several clinically relevant differences. For example, our finding of conserved CS in CIP^R^ mutants to GEN was previously reported in *E. coli* K12 (21) but not in MG1655 (20). In the CIP^R^ mutants we also observed conserved CR towards CHL, as reported in (20), but not in (22). We identified conserved CR of NIT^R^ mutants to AMX, and this was not reported in MG1655 (20). These observations underscore the importance of exploring collateral networks in multiple mutants of different clinical strain backgrounds and with different resistance mechanisms to assess their potential clinical application. Visual inspection of the data revealed a few clinically relevant examples of CS phenotypes that appeared independent of putative mechanism of resistance. We show that CIP^R^ *E. coli* strains displayed CS towards GEN, FOS, ETP and COL, and these phenotypes were conserved across multiple mechanisms of resistance. These results parallel those of a recent study on *P. aeruginosa* clinical isolates from cystic fibrosis patients, where CIP^R^ was associated with CS to GEN, FOS and COL. Taken together these data support the presence of general conserved collateral networks that may both affect the population dynamics of AMR during treatment and counter-select for resistance as recently indicated (30).

We assumed *a priori* that genetic background, AMR group, resistance mechanism, and the fitness cost of resistance could potentially affect the generality, sign and magnitude of collateral networks in clinical *E. coli* strains. Despite the fact that some collateral responses are conserved across different strains and mechanisms of resistance, our multivariate statistical approaches show overall that mechanism of resistance is the key predictor of CS and CR variability. This is primarily the case for efflux related mutations. However, mechanism of resistance also significantly contributed to the observed CS and CR variation in the MEC mutants where no efflux mutations were found. The presented data are consistent with earlier reports based on multiple AMR mutants derived from single strains with different resistance mechanisms towards specific antimicrobials (21, 22, 48). Our finding that genetic background did not significantly contribute to collateral responses is an important addition to these earlier studies. Finally, we found that the fitness cost of resistance also contributed significantly to the observed variation in CS and CR, despite some overlap in explanatory power due to the observed correlation between efflux-related mutations and reduced fitness. Taken together, our data and previous reports indicate that applied use of collateral networks in future treatment strategies may be dependent on rapid identification of specific resistance mechanisms. Moreover, clinical application of CS as a selection inversion strategy warrants further investigations to ideally explore CS in isogenic backgrounds, representing several diverse strains, with permutations of all known AMR-associated resistance traits. Such extensive studies would likely provide valuable information on the mechanisms of CS. Other confounding factors such as mobile genetic elements with heterogeneous resistance determinants should also be investigated as they would likely influence and reduce the predictability of collateral networks.

Selection inversion, as described by (20), depends on the cycling of drug pairs that display reciprocal CS. We did not observe reciprocal CS between any of the four drugs studied here that are widely used for treatment of urinary tract infections. However, we asked if modest reductions and increases in antimicrobial susceptibilities would affect the MSW (43) for the most prevalent CS and CR phenotypes. We subjected conserved CS and CR phenotypes to MPC assays and revealed that even a small 1.5-fold change in IC_90_ could equally alter the MPC, resulting in a shift of the MSW. These results suggest that antimicrobial treatment strategies informed by collateral networks could affect AMR evolutionary trajectories. Sequential treatment using drug pairs that display CR would, following resistance development, shift the MSW towards higher antimicrobial concentrations and increase the likelihood for resistance development to subsequent treatment options. Conversely, sequential treatment based on drug pairs that display CS can shift the MSW down and reduce the window of opportunity for high-level resistance development. This result suggests that the initial choice of antimicrobial may set the stage for later resistance development (Fig. 4). Based on our *in vitro* findings, TMP and NIT are attractive from a clinical perspective, as resistance to these resulted in few collateral responses, preserving the innate sensitivity to available secondary antimicrobials. However, MEC could be even more attractive, as CS largely dominates the observed collateral responses in MEC^R^ mutations. Additionally MEC^R^ isolates, especially those evolved *in vivo*, are associated with high cost of resistance (41). In contrast, CIP exposure was more likely to cause dramatic collateral responses that depended on the mechanism of resistance and could potentially negatively impact future therapeutic options. These observations align with antimicrobial treatment recommendations in Norway, where MEC, NIT and TMP are recommended for first line therapy of uncomplicated UTI, and CIP is reserved for otherwise complicated infections (49). Similarly, in the United States NIT, SXT (TMP-SMX) and MEC are recommended before fluoroquinolones such as CIP, ofloxacin, and levofloxacin (50).

**Figure 4:**
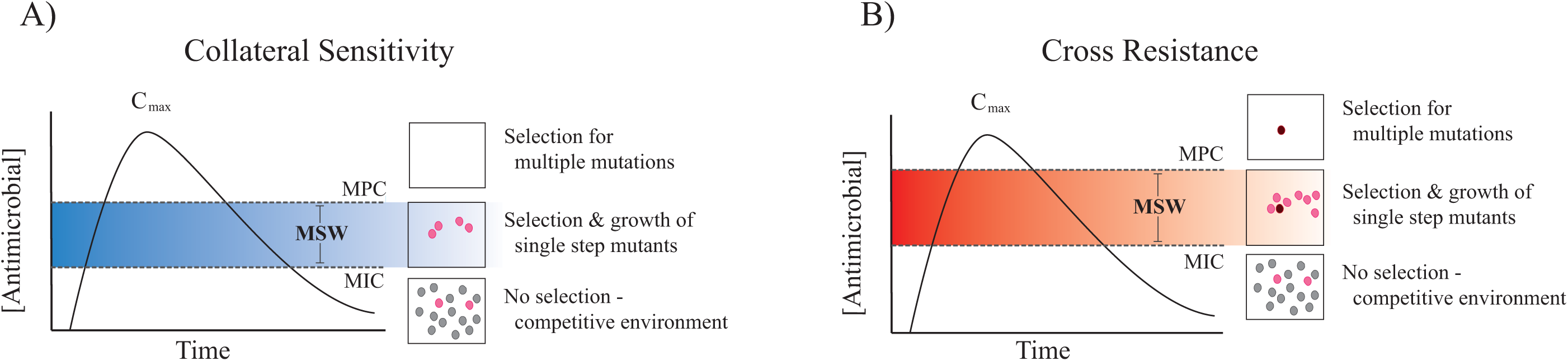
Graphical presentation of the potential effects of CS and CR on the MSW. Sequential drug administration informed by CS could potentially narrow or shift the MSW downwards in concentration space (left panel) whereas CR results in a widened or shifted upwards MSW (right panel). This would affect the probability of acquiring second step mutations leading to high-level resistance. Consequently, CS informed secondary therapies could reduce selection and thus propagation of first step mutants resulting in a reduced window of opportunity for second step mutations to occur. Dots represent bacteria resistant to a primary antibiotic (grey), spontaneous mutants with reduced susceptibility to a secondary drug (pink), or those with high-level resistance to the secondary drug (dark red).

Our conclusions are not without limitations. First, we acknowledge that including more clinical isolates from different infection foci, more diverse genetic backgrounds, as well as other selective agents, could change the outcome of our statistical analyses. This would allow increased sensitivity for the assessment of the different factors controlling collateral responses. A more targeted approach to assess the impact of specific resistance mechanisms on CS and CR across genetically diverse clinical strains is lacking in the field. Our analyses suggest that the fitness cost of resistance explains some variability in the collateral networks reported here. While it is known that growth rates affect antibiotic action (51), the underlying mechanism is currently unknown. It is also unclear if collateral networks will be perturbed by compensatory evolution, which eliminates the fitness costs of primary resistance (52-54). Finally, this and previous studies focus on AMR development to a single drug and there is a complete lack of data on how multidrug resistance, including resistance genes on mobile genetic elements, will affect collateral networks. We are currently investigating these and other questions that will aid in our understanding of collateral networks and their potential therapeutic application.

## Methods

### Bacterial strains

We used ten clinical urinary tract infection isolates of *E. coli* from the ECO-SENS collections (55, 56) originating from countries across Europe between 2000 and 2008 (Table 1). The isolates were chosen to represent pan-susceptible strains with diverse genetic backgrounds and were reported plasmid-free (31). Subsequent analysis based on whole-genome sequencing discovered two changes to previously reported ST’s and the presence of plasmid replicons in three strains (**Table S5E**). *E. coli* ATCC 25922 was used for reference and quality control purposes. For general growth, bacterial strains were grown in either Miller Difco Luria-Bertani (LB) broth (Becton, Dickinson and Co., Sparks MD, USA) or on LB agar; LB broth supplemented with select agar (Sigma-Aldrich) at 15 g/L. All strains were incubated at 37°C.

### Selection of AMR mutants

Single AMR mutants were selected at MICs above the European Committee on Antimicrobial Susceptibility Testing (EUCAST) clinical breakpoints (57) for CIP, NIT, TMP and MEC (Table 1). Resistant mutants were selected on Mueller Hinton II agar (MHA-SA; Sigma-Aldrich) for CIP, NIT and TMP using a step-wise static selection. MEC^R^ mutants were selected as single-step mutants on LB agar. For more details, see **Text S1**. AMR mutants were confirmed as *E. coli* using matrix-assisted laser desorption ionization time-of-flight (MALDI-TOF) analysis with MALDI BioTyper software (Bruker, MA, USA).

### Antimicrobial susceptibility testing

AMR mutants were screened for resistance above EUCAST breakpoints (**Table S1**, (57)) with gradient diffusion strips following manufacturers guidelines (Liofilchem, Italy), on Mueller Hinton II agar (MHA-BD; Becton, Dickinson and Company) after 18 hours incubation. Plates with insufficient growth were incubated an additional 24 hours.

Collateral changes to 16 antimicrobials (Table 2) were determined by IC_90_ testing (20), with some modifications. Standard 2-fold concentrations and median values between them were used as a “1.5-fold” testing scale. IC_90_ values were read as the first concentration tested that resulted in ≥ 90% inhibition of growth (optical density at 600nm, OD_600_) following 18 hours incubation at 700 rpm (3 mm stroke) in Mueller Hinton Broth (MHB, Becton, Dickinson and Company). Percent inhibition was calculated as previously described (20). IC_90_ results were determined in at least three biological replicates on separate days always including the control strain ATCC 25922. The final result reflects the average of a minimum of three replicates that met quality control standards (**Text S1**). Fold change in IC_90_ was calculated as the ratio between the AMR mutant and its respective WT.

Dose response curves were generated with averages of OD_600_ values (background subtracted) for each concentration tested during the IC_90_ experiments. Averages were plotted for AMR mutants and respective WT strains.

### Mutation prevention concentration testing

CS and CR trends to single drugs present in ≥ 50% of the isolates (CS_50_ or CR_50_), were confirmed by determining the mutation prevention concentrations (MPCs), essentially as described previously (58). Briefly, 10 mL aliquots of an overnight culture were pelleted and re-suspended in 1 mL MHB, estimated to contain ≥ 10^10^ CFU (actual values were 1.4 x 10^10^ - 7 x 10^10^ CFU). The inoculum was split and spread onto four large (14 cm diameter) MHA-SA agar plates for each antimicrobial concentration tested (4 - 6 concentrations of a 2-fold dilution series). The MPC was the lowest concentration with no visible growth after 48 h (**Text S1**).AMR mutants and WTs were tested in parallel and the results represent the average of a minimum of two biological replicates.

### Growth rate measurements

Growth curves of WT and AMR mutants were generated in a Versamax plate reader (Molecular Devices Corporation, California, USA) with constant shaking overnight. The OD_600_ was measured every 10 minutes and growth rates were estimated using GrowthRates v.2.1 software (59) (**Text S1**).

### Identification of genetic AMR determinants

Genomic DNA was isolated using the GenElute Bacterial Genomic DNA kit (Sigma-Aldrich) following guidelines for Gram-positive DNA extraction. Purity and quantification of genomic DNA was determined with Nanodrop (Thermo Scientific) and Qubit High Sensitivity DNA assay (Life Technologies), respectively and used to prepare libraries using the DNA Ultra II Library Preparation Kit (New England Biolabs, E7645) according to manufacturers description (**Text S1**). Libraries were then quantified by Qubit High Sensitivity DNA assay and distributions assessed by Bioanalyser DNA 1000 Chip (Agilent, 5067-1504) before normalizing and pooling. The final library pool was sequenced on the MiSeq (Illumina, San Diego) using 250 bp paired end reads and V2 chemistry.

WT genomes were assembled as described in **Text S1**. WT genomes were annotated with Rapid Annotation using Subsystem Technology server (RAST, version 2.0) for *E. coli* (60). SeqMan NGen (DNASTAR, Madison, WI) was used for comparative analysis of raw AMR mutant Illumina reads, using standard settings. AMR mutant reads were aligned to the corresponding annotated WT genome assembly. Reported SNPs had ≥ 10x coverage depth and ≥ 90% variant base calls. SNPs present in the WT assembly or in at least two AMR mutants of the same strain background were excluded. Genetic deletions and rearrangements were identified in the structural variation report generated in SeqMan Pro (DNASTAR) and were manually inspected and annotated using Gene Construction Kit (Textco Biosoftware Inc., Raleigh, NC) and NCBI BLAST searches, respectively.

### Multivariate Statistical Analyses

The fold changes of mean IC_90_ values (collateral responses) were determined relative to the parental WT strain and log transformed. Statistical analyses were performed on the complete data set, as well as a subset of the data excluding five antimicrobials, those to which primary AMR was evolved (CIP,MEC, NIT, and TMP) and SXT. To estimate and test the effects of strain background, AMR group, AMR mechanism, growth rate and relative fitness we relied on multivariate modeling, via redundancy analysis, to address the co-variation in IC_90_ across antimicrobials. The linear constraint scores were plotted for each AMR mutant. The response variables were overlaid with independent scaling to illustrate the direction of “steepest ascent” (increasing CR) from the origin for each antimicrobial. The significance of multivariate models and of their factors was assessed by permutation tests (1000 permutations) where p < 0.05 was considered significant. These analyses were done in R (61) using the Vegan work package (62).

### Data availability

Whole-genome sequencing data are available within the NCBI BioProject PRJNA419689. All other relevant data are available within this article, its Supplementary Information, or from the corresponding author upon request.

## Acknowledgments

We thank Tony Brooks at UCL Genomics for expertise in genome sequencing, Ane L.G. Utnes for generation of NIT^R^ mutants, Maria Chiara Di Luca for analysis of the genomes for MLST and identification of plasmid replicons, and Søren Overballe-Petersen for preliminary processing of WT genomes. Funding for this project was provided through the Northern Norway Regional Health Authority and UiT - The Arctic University of Norway (Project SFP1292-16), and JPI-EC-AMR (Project 271176/H10).

## Author Contributions

P.J.J. and 0.S. conceived the project; N.L.P., E.G.A.F., J.K., and V.S. designed and performed experiments; R.P. designed and R.P and N.L.P. performed the multivariate statistical modeling; all authors analyzed, interpreted and discussed the data; and N.L.P., E.G.A.F., A.P.R., D.E.R., and P.J.J. wrote the manuscript with contributions from the other authors.

## Supplementary Information List

### Text S1

**Figure S1:** The distribution of average IC_90_ values for wild-type *Escherichia coli* clinical isolates.

**Figure S2:** Collateral changes in susceptibility of 40 AMR mutants to 16 antimicrobials.

**Figure S3:** Dose response curves of representative strain:drug combinations that demonstrate frequently observed collateral responses (CS_50_ and CR_50_).

**Figure S4:** Relative growth rate of AMR mutants compared to their respective WT ancestors.

**Figure S5:** Multivariate models displaying the contribution of individual and combinations of factors to collateral networks.

**Table S1:** Gradient strip diffusion MIC values of AMR mutants.

**Table S2:** Tabulation of collateral responses detected in 40 AMR mutants.

**Table S3:** Genomic analyses of whole genome sequencing data in WT strains and AMR mutants.

**Table S4:** Summary of the output from multivariate models.

